# The Spatial Patterns and Determinants of Cerebrospinal Fluid Circulation in the Human Brain

**DOI:** 10.1101/2023.08.13.553149

**Authors:** Arash Nazeri, Taher Dehkharghanian, Kevin E. Lindsay, Pamela LaMontagne, Joshua S. Shimony, Tammie L.S. Benzinger, Aristeidis Sotiras

## Abstract

The circulation of cerebrospinal fluid (CSF) is essential for maintaining brain homeostasis and clearance, and impairments in its flow can lead to various brain disorders. Recent studies have shown that CSF circulation can be interrogated using low b-value diffusion magnetic resonance imaging (*low-b* dMRI). Nevertheless, the spatial organization of intracranial CSF flow dynamics remains largely elusive. Here, we developed a whole-brain voxel-based analysis framework, termed CSF pseudo-diffusion spatial statistics (CΨSS), to examine CSF mean pseudo-diffusivity (MΨ), a measure of CSF flow magnitude derived from *low-b* dMRI. We showed that intracranial CSF MΨ demonstrates characteristic covariance patterns by employing seed-based correlation analysis. Importantly, we applied non-negative matrix factorization analysis to further elucidate the covariance patterns of CSF MΨ in a hypothesis-free, data-driven way. We identified distinct CSF spaces that consistently displayed unique pseudo-diffusion characteristics across multiple imaging datasets. Our study revealed that age, sex, brain atrophy, ventricular anatomy, and cerebral perfusion differentially influence MΨ across these CSF spaces. Notably, individuals with anomalous CSF flow patterns displayed incidental findings on multimodal neuroradiological examinations. Our work sets forth a new paradigm to study CSF flow, with potential applications in clinical settings.

## Introduction

The cerebrospinal fluid (CSF) circulates through a complex network of inter-connected basilar cisterns, ventricular system, and extra-axial CSF spaces, which are semi-compartmentalized by brain parenchyma and meningeal membranes(*1*–*3*). Cardiovascular pulsatility drives CSF flow and propels it along the perivascular spaces (*4*), where CSF is in continuous exchange with the interstitial fluid and plays a vital role in maintaining brain homeostasis and clearance system (*1*, *5*). Disruptions in CSF circulation can lead to hydrocephalus and intracranial pressure disorders (*6*, *7*). Moreover, abnormalities in CSF flow dynamics have been observed in the context of aging, neurodegenerative disorders, and small vessel disease (*1*).

Prior work has used invasive and non-invasive techniques to study CSF flow in the human brain. However, studies using imaging approaches that involve intrathecal injection of contrast agents have often been limited by the number of participants or inclusion of healthy individuals due to the invasive nature of the procedure (*8*). Additionally, while CSF flow imaging techniques, such as phase-contrast imaging (*9*, *10*) and spin-labeling MRI (*11*, *12*), are non-invasive, they cannot provide whole-brain maps of CSF flow in relatively short acquisition times. This limitation has resulted in studies that have primarily focused on flow patterns across specific CSF passageways (e.g., cerebral aqueduct, foramina of Monro, and craniocervical junction) (*12*). Due to these limitations, the spatial organization of intracranial CSF flow dynamics has remained largely elusive.

Understanding CSF flow dynamics is further hindered by its complex nature. CSF flow is driven in tandem by oscillatory changes in brain volume across the cardiac and respiratory cycles and the pumping effects of cerebral vasculature (*13*–*15*). Importantly, these effects are moderated by intracranial anatomy and compliance of the CSF spaces (*1*, *16*). Therefore, comprehensive multimodal imaging studies are required to elucidate the effects of demographics, intracranial anatomy, and physiologic drivers on CSF flow patterns.

In this study, we sought to gain insight into the spatial organization of intracranial CSF circulation and characterize its underlying structural and physiological underpinnings. To this end, we capitalized on recent advances in diffusion-weighted MRI at low b-values (*low-b* dMRI) to evaluate CSF flow in the human brain (*17*–*21*). *Low-b* dMRI is particularly useful in this regard because it provides whole-brain maps of CSF flow in a non-invasive fashion. Accordingly, we derived maps of CSF mean pseudo-diffusivity (MΨ) (i.e., a measure of CSF flow magnitude derived from *low-b* dMRI (*18*)) in a large multi-b-value cohort, part of the Open Access Series of Imaging Studies (OASIS). We examined CSF MΨ throughout the brain using a whole-brain voxel-based analysis framework we developed, termed CSF pseudo-diffusion spatial statistics (CΨSS). Employing a hypothesis-driven approach, we applied seed-based correlation analysis to determine if the covariance patterns of CSF MΨ are consistent with the expected intracranial CSF flow patterns. Next, we utilized non-negative matrix factorization (NMF) to further analyze the covariance structure of CSF MΨ and elucidate the spatial patterns of CSF circulation in a data-driven and hypothesis-free way. In multiple independent *low-b* dMRI datasets, we showed that these CSF circulation spatial patterns represented distinct pseudo-diffusion magnitude and flow directionality. Finally, by employing multiple neuroimaging modalities, we identified the major determinants of CSF MΨ and showed that individuals with aberrant CSF flow patterns tended to have structural incidental findings on their neuroradiological examinations.

## Results

### Description of the low b-value diffusion MRI datasets

The OASIS-3 dataset included cognitively unimpaired participants as well as individuals with varying degrees of cognitive decline (Table 1). The multi-b-value imaging subset (n=187) of the OASIS-3 dataset was used as the primary dataset for CSF seed-based correlation analysis and NMF. Single shell *low-b* dMRI data from another OASIS-3 imaging subset (OASIS – b100 cohort, n=103, b-value = 100 s/mm^2^), the Microstructure-Informed Connectomics (MICA – b300, 49 healthy participants, b-value = 300 s/mm^2^), and the Comprehensive Diffusion MRI Dataset (CDMD – b50, 26 healthy participants, b-value = 50 s/mm^2^) were used for replication analyses (Table 1).

**Table 1.**
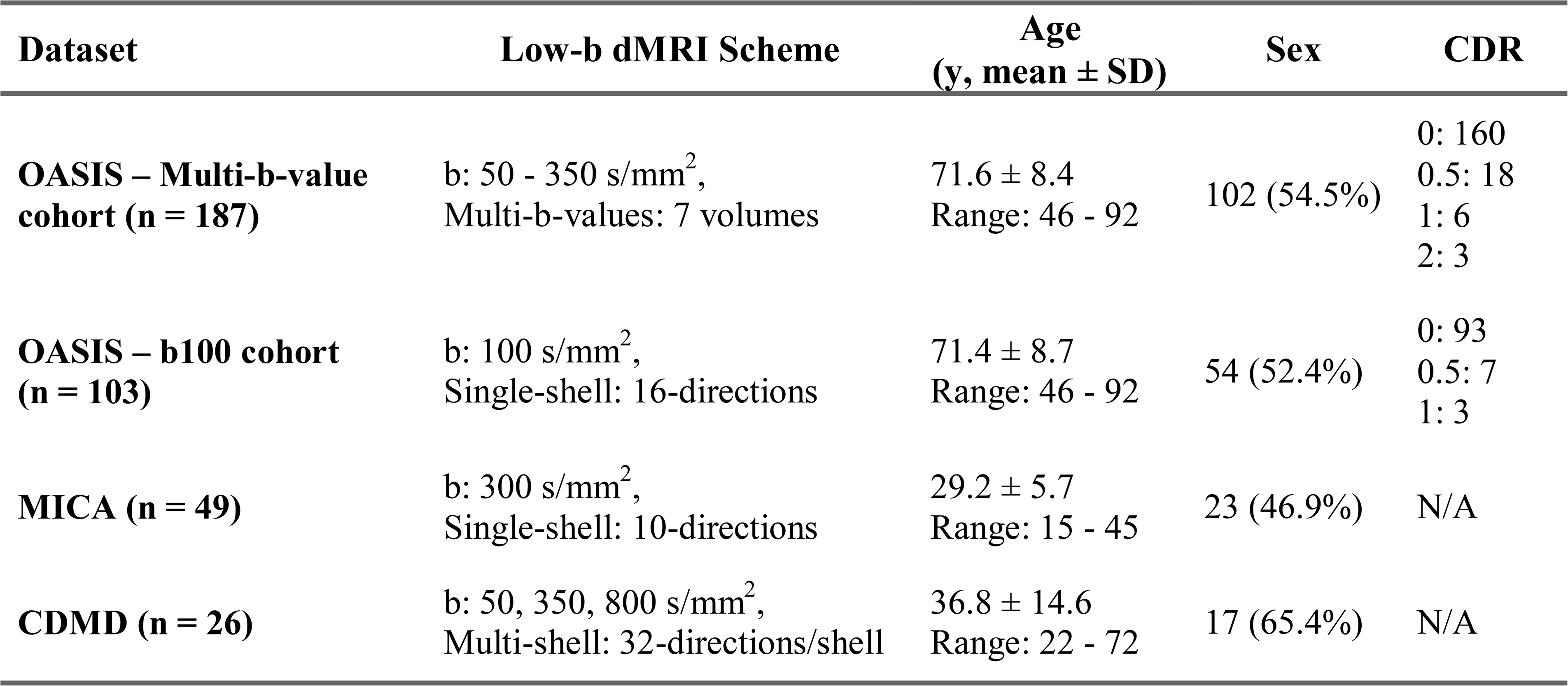
Demographic characteristics of the low b-value imaging datasets. CDR: Clinical Dementia Rating; dMRI: diffusion-weighted MRI; SD: standard deviation.

### CSF MΨ covariance aligns with the expected intracranial CSF flow patterns

A seed-based correlation analysis was conducted in the multi-b-value OASIS-3 subset to examine whether CSF MΨ covariance aligns with the expected interconnectivity of CSF flow throughout the brain. To prepare for voxel-wise analysis, CSF MΨ maps were aligned to a common space using CSF pseudo-diffusion spatial statistics (see Figure 1 for an overview of the CΨSS procedure). Five seed regions of interest (ROIs) were selected to explore CSF flow patterns in a hypothesis-driven way. These regions included the left and right Sylvian fissures to assess symmetry, the premedullary cistern to examine craniocervical flow, the cerebral aqueduct-fourth ventricle complex, and the foramina of Monro to analyze intraventricular flow (Figure 2).

**Figure 1.**
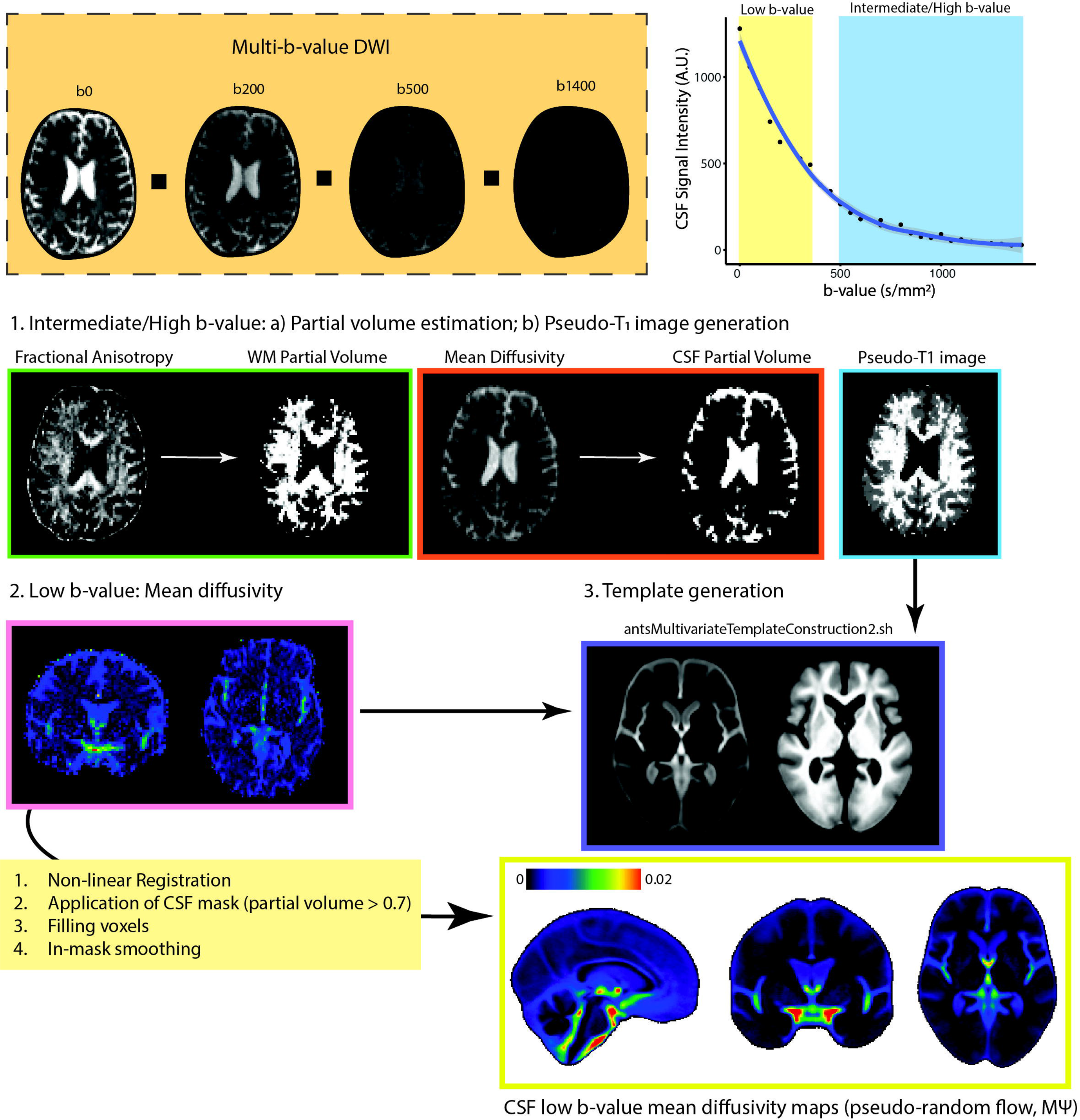
Overview of CSF Pseudo-diffusion Spatial Statistics (CΨSS). At low b-values, CSF retains it signal and signal changes are sensitive to incoherent flow. For CΨSS, the following steps were performed: (1) Diffusion tensor model was applied to intermediate/high b-value images to obtain FA and mean diffusivity. These were used for 2-tissue segmentation (Atropos, part of ANTs) to generate CSF partial volume maps and pseudo-T1-weighted images. (2) Low b-value diffusion-weighted images (multi-low b-value subset: b: 0-350 s/mm^2^ [7 non-b0 volumes]; b100 subset: b = 100 s/mm^2^, 16 directions, 3 b0 volumes) were used to generate low b-value MΨ maps, measuring pseudorandom flow magnitude. (3) MΨ and pseudo-T1 images were utilized to create study-specific templates with *antsMultivariateTemplateConstruction2.sh* (ANTs).Subsequently, MΨ and CSF partial volume maps were registered to the template space. A final CSF mask was generated by retaining voxels with CSF fraction > 0.7 in more than 65% subjects. Voxels with a CSF fraction < 0.7 were filled using surrounding voxel average. Spatial smoothing (3 mm FWHM) was applied within the CSF mask using 3dBlurInMask (part of AFNI). *Abbreviations:* CSF, cerebrospinal fluid; FA, fractional anisotropy; FWHM, full width half maximum; MΨ, mean pseudo-diffusivity.

**Figure 2.**
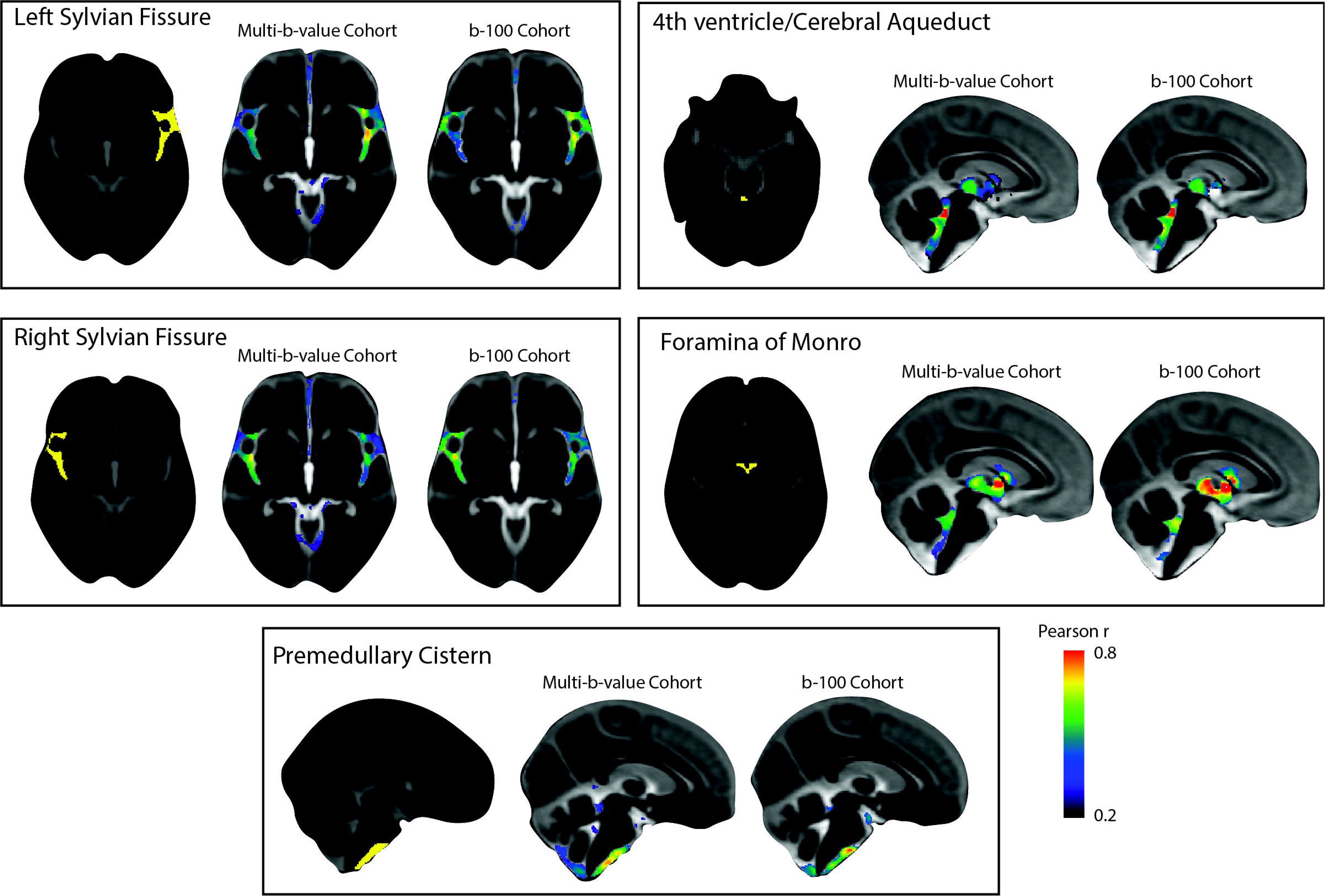
Patterns of CSF MΨ covariance. Hypothesis-driven seed-based correlation analysis was conducted to investigate CSF flow covariance patterns. Similar seed-based correlation maps were observed in both OASIS-3 imaging cohorts (multi-b-value and b100 imaging cohorts). CSF MΨ in the left and right Sylvian fissures displayed similar correlation patterns that demonstrated high bilateral symmetry. The correlation maps of the cerebral aqueduct and foramina of Monro MΨ were similar, reflecting the common pathway for trans-ventricular CSF flow that included the third and fourth ventricles. Premedullary CSF MΨ was positively correlated with CSF flow in the dorsal and ventral craniocervical junction columns extending to the cerebellopontine angles. *Abbreviations:* CSF, cerebrospinal fluid; MΨ, mean pseudo-diffusivity.

CSF MΨ in the left and right Sylvian fissures displayed similar correlation patterns that demonstrated high bilateral symmetry with higher correlation values observed in regions ipsilateral to the seed than contralateral ones. The correlation patterns included the perivascular CSF spaces surrounding the main arteries of the anterior circulation in the ipsilateral and contralateral Sylvian fissures and carotid cisterns, as well as the anterior interhemispheric subarachnoid space. The correlation maps of the cerebral aqueduct and foramina of Monro MΨ were similar, reflecting the common pathway for trans-ventricular CSF flow, including the third and fourth ventricles. Premedullary CSF MΨ was positively correlated with CSF flow in the dorsal and ventral craniocervical junction columns extending to the cerebellopontine angles. Similar seed-based correlation maps were also observed in the b100 imaging cohort (Figure 2) with Pearson r values between correlation maps generated from the multi-b-value and b100 cohorts ranging from 0.66 – 0.78.

### NMF reveals the organization of CSF flow within the ventricular system and basilar cisterns

Having demonstrated that the covariance of CSF MΨ is in accordance with the expected patterns of CSF flow, our next objective was to investigate the organization of intracranial CSF flow using an unbiased, data-driven approach. To achieve this, we applied NMF to the CSF MΨ maps from the OASIS-3 multi-b-value cohort. NMF is a multivariate pattern analysis technique, which has recently been adapted for use in neuroimaging (*22*–*24*) and allows for a parts-based representation of complex high dimensional datasets (*25*–*27*). We evaluated multiple NMF solutions with varying numbers of patterns (K = 2 to 25) to determine the most reproducible and reliable solution with the lowest reconstruction error. Split-half reproducibility analysis, as quantified using the Adjusted Rand Index (ARI) (*28*), demonstrated nonuniform reproducibility and reliability across NMF solutions (Figure 3A). Reproducibility (i.e., high mean ARI across bootstraps) peaked at the 9- and 10-pattern solutions (mean ARI: 0.81 [9-pattern] and 0.82 [10-pattern]). The highest reliability (i.e., high stability with low ARI standard deviation across bootstraps) was also observed for the 9-pattern (SD=0.017) and 10-pattern (SD=0.018) solutions (Figure 3A). The 10-pattern solution, which had a lower reconstruction error, was selected for the subsequent analyses (Figure 3B).

**Figure 3.**
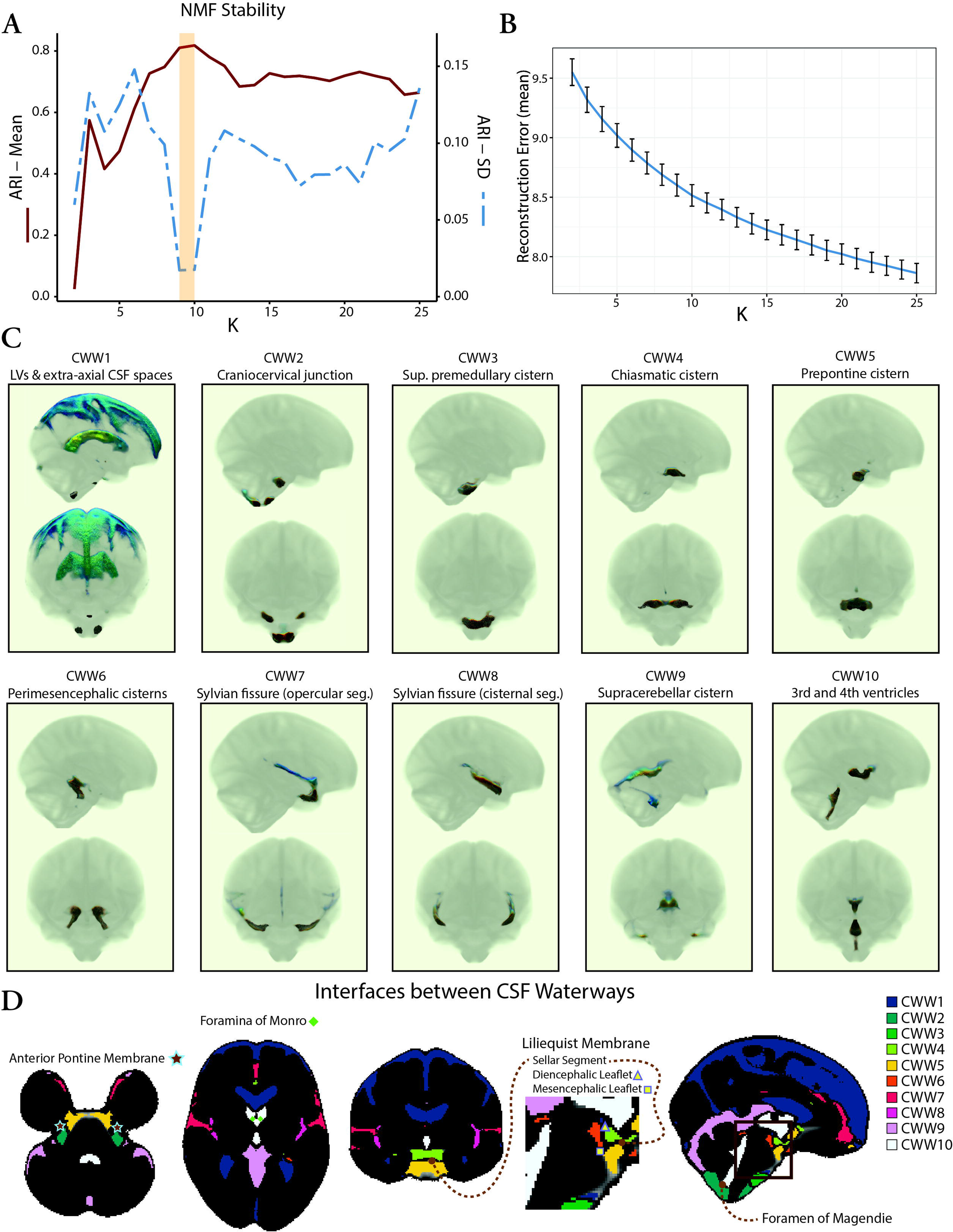
Coordinated CSF flow organization in CSF waterways (CWWs) uncovered by non-negative matrix factorization. (A) Evaluation of NMF solutions (K = 2 to 25) showed that the 9-pattern and 10-pattern solutions had the highest mean ARI and lowest ARI standard deviation, indicating highest reproducibility and reliability, respectively. (B) Reconstruction errors decreased with higher NMF solutions. The 10-pattern solution, with a lower reconstruction error, was selected for further analysis. (C) The CSF waterways (CWWs; the NMF patterns) were symmetrically organized within the ventricular system and basilar cisterns: CWW1, lateral ventricles and extra-axial subarachnoid spaces; CWW2, cisterna magna, cervicomedullary cistern, and cerebellopontine angles; CWW3, superior premedullary cistern; CWW4, chiasmatic cistern; CWW5, prepontine cistern; CWW6, perimesencephalic cisterns and choroid fissures; CWW7, superficial opercular compartments of the Sylvian fissure; CWW8, deep cisternal portions of the Sylvian fissure; CWW9, supracerebellar and velum interpositum cisterns; CWW10, trans-ventricular CSF pathway. (D) The CWWs separated by functional and structural boundaries. Trans-ventricular CWW10 was bounded by Monro and Magendie foramina. Interfaces of basilar cistern CWWs aligned with arachnoid membranes, including the Liliequist and anterior pontine membranes. *Abbreviations:* ARI, Adjusted Rand Index; CSF, cerebrospinal fluid; CWWs, CSF waterways, NMF, non-negative matrix factorization.

The CSF waterways (CWWs; the NMF patterns) were symmetrically organized within the ventricular system and basilar cisterns (Figure 3C). The ventricular system was subdivided into the lateral ventricle CWW1 and the trans-ventricular CWW10. CWW1 also included the extra-axial subarachnoid spaces, whereas the trans-ventricular CWW10 encompassed a small portion of the lateral ventricles adjacent to the foramina of Monro, the third and fourth ventricles, and the cerebral aqueduct. The Sylvian fissures were symmetrically divided into the deep cisternal (CWW8) and superficial opercular compartments (CWW7) (*29*). Six other CWWs corresponded to the basilar cisterns: (i) cisterna magna, cervicomedullary cistern, and cerebellopontine angles (CWW2); (ii) superior premedullary cistern (CWW3); (iii) chiasmatic cistern (CWW4); (iv) prepontine cistern (CWW5); (v) perimesencephalic cisterns and choroid fissures (CWW6); (vi) supracerebellar and velum interpositum cisterns (CWW9).

### CSF waterways displayed distinct pseudo-diffusion characteristics

To investigate whether the identified CWWs also corresponded to CSF spaces with distinct pseudo-diffusion characteristics (across the population), we extracted the mean CSF MΨ values from each CWW for every participant. The CWWs displayed noticeable differences in their CSF MΨ distributions in the OASIS multi-b-value cohort (Figure 4A). Specifically, the superior premedullary cistern (CWW3) exhibited the highest MΨ followed by the craniocervical junction (CWW2), the chiasmatic cistern (CWW4), and the prepontine cistern (CWW5). Conversely, the extra-axial/lateral ventricular CSF (CWW1) displayed the lowest MΨ. Among the basilar cisterns, the supracerebellar (CWW9) and perimesencephalic cisterns (CWW6) exhibited the lowest MΨ. Furthermore, the deep cisternal compartment of the Sylvian fissure (CWW8) consistently exhibited higher MΨ than the superficial opercular Sylvian fissure (CWW7). Similar mean MΨ distributions were observed across CWWs in three separate single-shell diffusion-weighted MRI datasets (Figure 4A) with different low b-values (OASIS – b100, MICA b300, and CDMD – b50). Notably, the MΨ values showed the lowest dynamic range in MICA b300 cohort.

**Figure 4.**
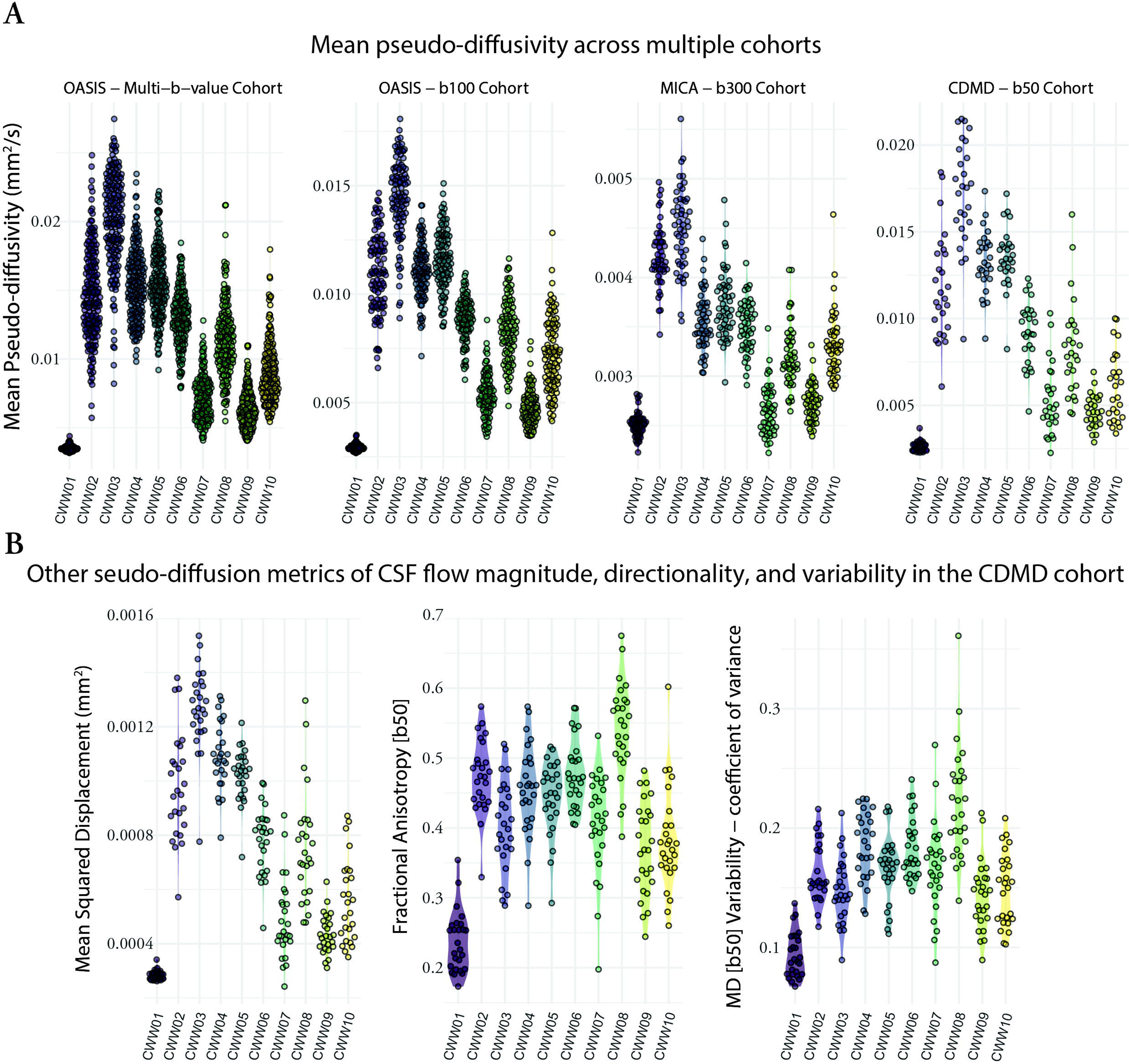
Pseudo-diffusion characteristics of the CSF waterways (CWWs). (A) The CWWs displayed distinct CSF MΨ distributions. Similar patterns were observed across CWWs in three separate single-shell diffusion-weighted MRI datasets. (B) Densely sampled diffusion MRI data from the CDMD showed that CWWs represent CSF spaces with distinct CSF incoherent flow magnitude (MSD), directionality (FA), and MΨ variability. *Abbreviations:* CSF, cerebrospinal fluid; CWWs, CSF waterways; FA, fractional anisotropy, MΨ, mean pseudo-diffusivity; MSD, mean square displacement.

We leveraged the densely sampled diffusion MRI data from the CDMD to explore additional pseudo-diffusion characteristics of CSF (Figure 4B). First, we calculated mean square displacement (MSD) values derived from the multi-shell *low-b* dMRI (b-values: 50 s/mm^2^, 350 s/mm^2^, 800 s/mm^2^, 32 directions for each shell). MSD is a measure of the distance the water molecules travel during the diffusion time and is closely related to mean diffusivity. As expected, multi-shell MSD values were highly correlated with single-shell MΨ values across the CWWs (Pearson r: 0.83 – 0.99; *p* < 0.001), indicating that the use of single-shell *low-b* dMRI is sufficient for evaluation of pseudo-diffusion magnitude. Next, we assessed the directionality of incoherent CSF flow in different CWWs by calculating fractional anisotropy (FA) at b=50 s/mm^2^. Apart from the extra-axial/lateral ventricular CSF CWW1 that showed relatively isotropic pseudo-diffusion (FA: 0.17 – 0.35), other CWWs showed either high (CWW8) or intermediate anisotropy (Figure 4B). Unlike MSD, the correlation between CSF FA and MΨ was highly variable across different CWWs (Figure S1). Finally, to assess the variability of MΨ, coefficient of variance of MΨ values was calculated across 100 sub-samples of b = 50 s/mm^2^ dMRI data (each subsample consisting of 12 volumes). We observed the lowest MΨ variability in CWW1, followed by CWW9 and CWW10, while CWW8 demonstrated the highest MΨ variability. Taken together, these findings suggest that CWWs represent CSF spaces with distinct CSF pseudo-diffusivity, flow directionality, and flow variability.

### Demographics, brain anatomy, and perfusion influence CSF pseudo-diffusion in a regionally specific way

The data-driven NMF approach identified patterns of CSF MΨ covariance, but did not include information regarding participant demographics, intracranial anatomy, or cerebral perfusion. Accordingly, in both OASIS-3 cohorts, we examined how age, sex, brain atrophy, ventricular anatomy, and brain perfusion affect CSF MΨ (Figure 5). Advancing age was associated with higher MΨ values in the Sylvian fissure CWWs (CWW7 and CWW8) and the chiasmatic cistern (CWW4), which are adjacent to the middle cerebral artery branches and circle of Willis, respectively. Women showed higher CSF MΨ values in the caudal basilar cisterns (CWW2 and CWW3).

**Figure 5.**
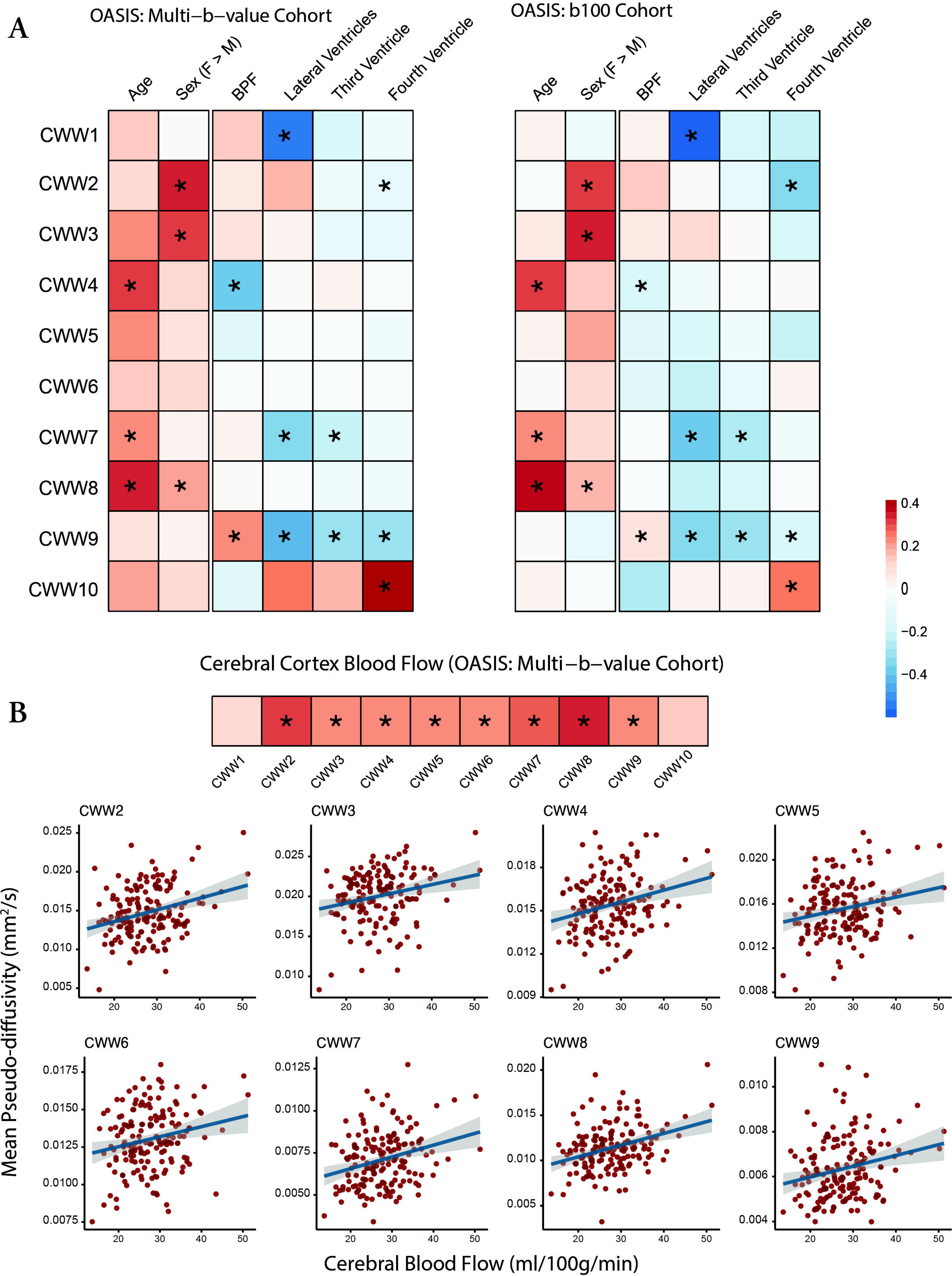
Effects of demographics, brain anatomy, and perfusion on CWW mean pseudo-diffusivity. (A) Effects of age, sex, and brain on CWW MΨ were estimated using the following model: CWW_MΨ ∼ age + sex + BPF. Effects of ventricular volumes and CBF were estimated while accounting for the effects of age, sex, and BPF using the following model: CWW_MΨ ∼ age + sex + BPF + [normalized ventricular volume]. Asterisks (*) indicate significant effects in the aggregate model of both OASIS-3 cohorts (FDR-corrected *p* < 0.05). (B) Gray matter CBF values were derived from pulsed arterial spin labeling (PASL). Effects of CBF were estimated while accounting for the effects of age, sex, and BPF using the following model: CWW_MΨ ∼ age + sex + BPF + CBF. Asterisks (*) indicate significant effects (FDR-corrected *p* < 0.05). *Abbreviations:* BPF, brain parenchymal fraction; CBF, cerebral blood flow, CWWs, CSF waterways; FDR, false discovery rate; MΨ, mean pseudo-diffusivity.

Brain atrophy (i.e., lower brain parenchymal fraction) and larger ventricle volumes were associated with lower supracerebellar cistern CWW9 MΨ. Larger lateral ventricle volume was also associated with lower MΨ in the extra-axial/lateral ventricular CSF (CWW1) and superficial opercular Sylvian fissure (CWW7). Larger fourth ventricle volume was associated with higher transventricular CWW10 MΨ and lower craniocervical junction CWW2 MΨ.

In the multi-b-value OASIS cohort, we used the available arterial spin labeling (ASL) imaging data to investigate the association between cerebral perfusion and CSF flow within CWWs. Except for the ventricular CWWs (CWW1 and CWW10), higher cerebral blood flow was associated with a higher CSF MΨ (Figure 5B). Strongest associations were observed in the Sylvian fissure CWWs (CWW7 and CWW8) and craniocervical junction CWW2. Overall, these findings highlight the regional specificity of the effects of demographics, intracranial anatomy, and cerebral perfusion on CSF flow.

### Aberrant CSF flow patterns were accompanied by incidental findings

In the OASIS study, individuals were considered outliers if they exhibited (i) a single CWW with an MΨ above or below 3 standard deviations from the mean or (ii) two or more CWWs with MΨ values above or below 2 standard deviations from the mean. Out of thirty-eight participants with outlier CWW MΨ values (multi-b-value cohort: 23; b100 cohort: 15), five individuals had clinically significant findings (Figure 6) including: bilateral subdural effusions (IF001); fourth ventricle mass (IF002); multiple intracranial aneurysms (IF003); retinal detachment and new cerebellar and cortical cystic lesions (IF004); cerebellar arteriovenous malformation (IF005). Craniocervical incidental findings were observed in two other participants with low CSF flow at the craniocervical junction (CWW2) or caudal basilar cisterns (CWW3 and CWW5): mega cisterna magna (IF006) and tonsillar ectopia (IF007).

**Figure 6.**
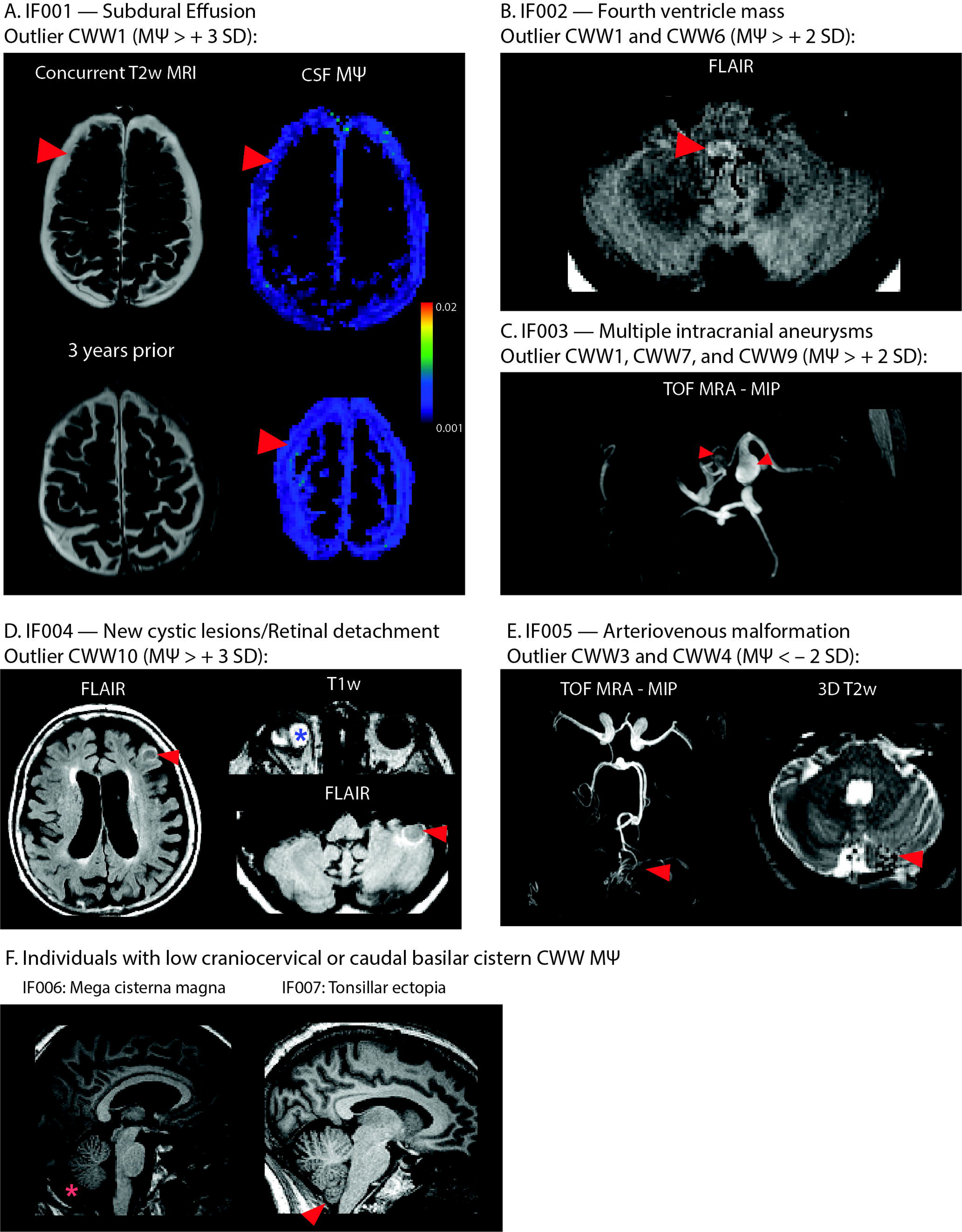
Incidental findings associated with aberrant CSF flow patterns. (A) IF001 exhibited high MΨ values in the subdural fluid collections, particularly adjacent to the arachnoid matter interface. (B) MRI images of IF002 revealed a FLAIR-hyperintense nodular lesion within the caudal fourth ventricle, narrowing the foramen of Magendie. (C) Time-of-flight MRA showed multiple intracranial aneurysms in IF003. (D) IF004 exhibited retinal detachment in the right eye, alongside newly identified cerebellar and cortical cystic lesions. (E) Time-of-flight MRA of IF005 showed arteriovenous malformation arising from the left superior cerebellar artery. Flow voids were evident in the left cerebellar hemisphere. (F) Two individuals with low CSF flow at craniocervical junction or caudal basilar cisterns showed craniocervical incidental findings: mega cisterna magna (IF006) and tonsillar ectopia (IF007). *Abbreviations:* CWWs, CSF waterways; FLAIR, fluid-attenuated inversion recovery; MΨ, mean pseudo-diffusivity; MRA, magnetic resonance angiography.

## Discussion

In this work, we demonstrated the intricate interconnectivity of intracranial CSF circulation using a novel voxelwise CSF flow mapping framework and large whole-brain *low-b* dMRI datasets. We identified distinct spatial patterns of CSF flow using an unsupervised multivariate pattern analysis approach. We showed that the identified patterns correspond to distinct CSF pseudo-diffusion characteristics and are differentially impacted by age, sex, ventricular anatomy, and brain perfusion. Our results were replicated in independent datasets with different *low-b* dMRI protocols. Finally, we showed that individuals with aberrant CSF flow patterns tend to exhibit incidental neuroradiological findings.

Consistent with prior reports (*17*–*19*, *21*), we demonstrate that *low-b* dMRI can rapidly probe CSF flow throughout the entire brain within a clinically feasible timeframe (1-3 minutes). Additionally, our study revealed that variations of the MΨ values across different CSF spaces are consistent across a range of imaging protocols, including those with different b-values (ranging from 50 s/mm^2^ – 350 s/mm^2^), single-shell and multi-b-value imaging schemes, various spatial resolutions, and MRI hardware. Further, we showed that other characteristics of CSF flow such as anisotropy and pseudo-diffusion variability can also be assessed using *low-b* dMRI with higher angular resolution. These findings suggest that *low-b* dMRI offers a practical, reliable, and versatile method for investigating CSF flow and may facilitate diagnosis and monitoring of brain disorders associated with altered CSF dynamics.

Our hypothesis-driven and data-driven results illustrate the organization of intracranial CSF flow into symmetric anatomical and functional units. The symmetry of CWWs with non-communicated compartments such as the lateral ventricles, extra-axial subarachnoid spaces, and Sylvian fissures suggests that CSF flow dynamics within these CSF spaces are driven and constrained by similar underlying mechanisms with left-right symmetry. In addition, the CWWs were segregated by functional and structural boundaries (Figure 3D). The trans-ventricular CWW10 was bordered by the foramina of Monro and Magendie, and was further demarcated by fine parenchymal structures such as the lamina terminalis (separating the third ventricle from the lamina terminalis cistern [CWW5]). In some other instances, the interfaces between basilar cistern CWWs aligned with the arachnoid membranes (*3*, *30*). The interface separating the chiasmatic cistern (CWW4) and prepontine cisterns (CWW5) approximated the sellar segment of the Liliequist membrane. The interface separating the interpeduncular cistern (CWW6) from the prepontine (CWW5) and chiasmatic (CWW4) cisterns followed the mesencephalic and diencephalic leaflets of the Liliequist membrane, respectively. The correspondence between the interfaces of basilar cistern CWWs and arachnoid membranes, such as the Liliequist membrane (*3*), underscores the influence of arachnoid matter’s delicate anatomy on shaping intracranial CSF flow patterns.

We found that the ventricular system could be functionally divided into low isotropic CSF flow in the lateral ventricles (CWW1) and transventricular anisotropic and variable CSF flow (CWW10) encompassing the frontal horns of the lateral ventricles adjacent to the foramina of Monro, third ventricle, cerebral aqueduct, and the fourth ventricle. Moreover, these intraventricular CSF flow patterns were influenced by distinct intracranial anatomy. In the lateral ventricles, larger volumes were associated with lower CSF MΨ, suggesting higher ventricular compliance results in dampened intraventricular pressure variation and CSF flow across the cardiac cycle. In contrast, our results indicate that larger fourth ventricular volumes are associated with higher transventricular CSF MΨ, which is in concordance with the diffuser/nozzle pump model, which posits that fourth ventricular morphology affects CSF flow in the third ventricle–aqueduct–fourth ventricle complex (*31*). Of note, these relationships, primarily reflecting normal variations in ventricle volumes, could potentially undergo changes in the setting of disease conditions such as hydrocephalus.

The association between cerebral blood flow and CSF MΨ in the caudal basilar cisterns likely reflects the coupling between the fluctuations in the cerebral blood volume and intracranial CSF volume throughout the cardiac cycle, as asserted by the Monro-Kellie doctrine (*32*). Our results also show that higher cerebral blood flow is associated with increased MΨ in the peri-arterial CSF spaces. The peri-arterial CSF flow is propelled by arterial pulsations (*13*, *14*), which could be amplified by higher cerebral blood flow. The observed age-related increase in periarterial CSF MΨ aligns with recent findings of elevated peri-arterial CSF flow and pulsatility in aging subjects (*21*). This rise in peri-arterial CSF flow is likely driven by increased intracranial arterial pulsatility with aging (*33*).

By reviewing multimodal images of the OASIS study participants with aberrant CSF flow dynamics, we were able to identify various incidental findings ranging from vascular anomalies to mass lesions, craniocervical junction findings, and intracranial fluid collections. In some instances, a direct causal link could be established between the observed CSF flow abnormality and the incidental findings. For instance, high CWW1 MΨ associated with subdural effusions suggests higher flow within the subdural collections compared to the extra-axial subarachnoid space, particularly adjacent to the bridging veins and arachnoid membrane. Taken together, our results highlight the importance of considering CSF dynamics in the context of a broad range of brain disorders and suggest that our approach may have implications for the diagnosis and treatment of a wide range of brain disorders.

There are some limitations that need to be considered. While in this study we capitalized on a large number of *low-b* dMRI images to evaluate the patterns of CSF flow and the factors influencing them, the *low-b* dMRI used in these datasets were acquired as a part of multi b-value datasets. Future studies can exploit the inherent high signal-to-noise ratio of *low-b* dMRI images and the possibility to acquire images at short echo times to collect dedicated *low-b* dMRI for high spatial resolution CSF flow imaging with minimal image distortion or blurring. Such high-fidelity, high-spatial-resolution *low-b* dMRI is particularly crucial for evaluating relatively narrow sulcal subarachnoid spaces and brain areas adjacent to air-tissue interfaces. Moreover, in this study, the *low-b* dMRI images were not cardiac gated. Recent research has shown that gated *low-b* dMRI allows for characterizing the dynamics of CSF flow over the cardiac cycle (*21*). Finally, while our results provide valuable insights into CSF flow under normal conditions, future studies are needed to apply this technique to investigate alterations in CSF flow in various disease conditions.

In conclusion, our work sets forth a new paradigm to study CSF flow by capitalizing on large sample size *low-b* dMRI datasets and unsupervised machine learning. Our study provides insights in the CSF flow organization, revealing distinct CSF waterways, which exhibit distinct CSF pseudo-diffusion characteristics and are differentially impacted by age, sex, ventricular anatomy, and brain perfusion. The data-driven CWW-based atlas of the intracranial CSF flow offers a systematic, parts-based approach to examining intracranial CSF flow patterns under physiological conditions and their abnormalities in the setting of various brain CSF disorders such as hydrocephalus, idiopathic intracranial hypertension, or Chiari 1 malformation.

## Methods

### Participants

#### Open Access Series of Imaging Studies

OASIS-3 (Open Access Series of Imaging Studies: http://www.oasis-brains.org/) is a retrospective compilation of clinical and imaging data from >1,300 participants that were collected across several studies through the Charles F. and Joanne Knight Alzheimer Disease Research Center (Knight ADRC: https://knightadrc.wustl.edu/) at Washington University in St. Louis (*34*). Participants include cognitively unimpaired adults and individuals at various stages of cognitive decline. This study utilized *low-b* dMRI data from two imaging cohorts within OASIS-3 that employed distinct imaging protocols and MRI scanners (Table 1): i) multi-low b-value subset: n=187 (after exclusion of 10 participants due to motion artifacts), 103 female, average age: 71.5 years [range: 46.2-92.2 years], 163 cognitively normal (Clinical Dementia Rating [CDR]: 0) and 18 individuals with very mild cognitive impairment (CDR: 0.5); ii) b100 subset: n=103 (after exclusion of a single participant due to motion artifact), 54 female, average age: 71.4 years [range: 46-91.6 years], 92 cognitively normal (CDR: 0) and 8 individuals with very mild cognitive impairment (CDR: 0.5).

#### The Microstructure-Informed Connectomics dataset

The Microstructure-Informed Connectomics (MICA) dataset (*35*) provides raw neuroimaging data collected from 50 healthy participants (Table 1). The imaging dataset is available on the Canadian Open Neuroscience Platform’s data portal (https://portal.conp.ca).

#### The Comprehensive Diffusion MRI Dataset

The Comprehensive Diffusion MRI Dataset (CDMD) included imaging data from 26 healthy participants (Table 1). The imaging dataset is available as a figshare collection (https://doi.org/10.6084/m9.figshare.c.5315474.v1).

### MRI acquisition

#### Open Access Series of Imaging Studies

The imaging cohorts in the OASIS-3 dataset were defined based on their dMRI imaging schemes. For the subset of OASIS-3 participants in the multi-low b-value group, brain MRI scans were conducted using a 3 T BioGraph mMR scanner (Siemens, Erlangen, Germany). Axial diffusion-weighed pulsed-gradient spin-echo echoplanar images (EPI) were acquired using a multi-low b-value protocol with the following imaging parameters: TE, 86 ms; TR, 10,300 ms; voxel size, 2 × 2 × 2 mm^3^; field-of-view, 224 × 224 mm^2^; slice number, 80. Each diffusion gradient had a unique b-value ranging from 0 to 1400 s/mm2, with a total of 26 volumes. For the subset of OASIS-3 participants in the b100 group, whole-brain diffusion-weighted EPI images were obtained using a 3 T Magnetom Vida scanner (Siemens, Erlangen, Germany) equipped with a 64-channel head coil with the following imaging parameters: TE, 79 ms; TR, 5,800 ms; voxel size, 2 × 2 × 2 mm^3^; field-of-view, 220 × 220 mm^2^; slice number, 80; multi-band factor, 2. Diffusion-weighted images were acquired in the axial plane along 66 gradient directions, with b-values of 100 s/mm^2^ (16 directions), 250 s/mm^2^ (10 directions), 500 s/mm^2^ (12 directions), 1,000 s/mm^2^ (12 directions), 1,500 s/mm^2^ (10 directions), 2,000 s/mm^2^ (6 directions), and 3 b0 volumes. Additionally, pulsed arterial spin labeling (PASL) imaging data was available for a subset of participants (n=164) (*34*). Axial 2D PASL images were acquired with proximal inversion with control of off-resonance effects (PICORE) tagging method and the following imaging parameters: TR, 3400 ms; TE, 13 ms; TI_1_,700; TI_2_, 1900 ms; in-plane resolution, 4 ×LJ4LJmm^2^; slice thickness,LJ5LJmm; 9 slices; 52 label/control pairs with an M0 reference image.

#### The Microstructure-Informed Connectomics dataset

In the MICA study(*35*), brain MRI images were acquired with a 3LJT Magnetom Prisma-Fit scanner (Siemens, Erlangen, Germany) equipped with a 64-channel head coil. Multi-shell diffusion MRI data were acquired using a diffusion-weighed pulsed-gradient spin-echo EPI sequence with the following parameters: 10 diffusion weighting directions at b=300 s/mm^2^ (only b = 300 s/mm^2^ and b0 images were used for this study), TE, 64.40 ms; TR, 3500 ms; voxel size, 1.6 × 1.6 × 1.6 mm^3^, field-of-view,LJ224LJ×LJ224LJmm^2^; multi-band factor, 3. To correct for susceptibility induced image-distortions of, b0 images were also acquired in reverse phase encoding direction.

#### The Comprehensive Diffusion MRI Dataset

In the CDMD study, all data were acquired on the 3LJT Magnetom Connectome MRI scanner (Siemens, Erlangen, Germany) equipped with a maximum gradient strength of 300 mT/m and a custom-built 64-channel phased array head coil(*36*). Multi-shell diffusion MRI data were acquired using a sagittal diffusion-weighed pulsed-gradient spin-echo EPI sequence with the following parameters: TE, 77 ms; TR, 3800 ms; voxel sizeLJ=LJ2LJ×LJ2LJ×LJ2 mm^3^, field-of-view, 216LJ×LJ216LJmm, multi-band factor, 2. Diffusion-weighted images acquired at three different b-values were used in this study (b = 50, 350, 800 s/mm^2^; acquired along 32 diffusion encoding directions uniformly distributed on a sphere). Five b0 image volumes with reversed phase-encoding direction were acquired to correct for susceptibility-induced image distortions.

### CSF pseudo-diffusion spatial statistics (CΨSS)

After corrections for Gibbs ring artifact (*mrdegibbs* tool, part of MRtrix3) (*37*, *38*) and eddy current-induced distortions (*eddy_correct* tool with spline interpolation, part of FSL) (*39*), skull stripping was performed (Brain Extraction Tool [BET], part of FSL) (*40*). The diffusion tensor model was fitted to the low-b-value diffusion-weighted images (multi-low b-value subset: b: 0-350 s/mm^2^ [7 non-b0 volumes]; b100 subset: b = 100 s/mm^2^, 16 directions, 3 b0 volumes) to generate low b-value MΨ maps as a measure of pseudorandom flow magnitude. The diffusion tensor model was also fitted into the intermediate/high b-value images in both datasets (multi-low b-value subset, b: 500 – 1,400 s/mm^2^; b100 subset, b: 500 – 2,000 s/mm^2^). The resulting FA and mean diffusivity images were fed into a 2-tissue segmentation algorithm (*Atropos*, part of ANTs; http://stnava.github.io/ANTs/) to generate white matter and CSF partial volume maps(*41*). Similar to gray matter-based spatial statistics(*42*, *43*), partial volume maps were used to generate pseudo-T_1_-weighted images. MΨ and pseudo-T_1_ images were used to create study-specific templates using the *antsMultivariateTemplateConstruction2.sh* workflow (Figure 1). MΨ and CSF partial volume maps were subsequently registered to the template space. The final mask was generated by keeping voxels with a CSF fraction greater than 0.7 in more than 65% of the subjects. Voxels with CSF fraction < 0.7 were then filled with the average of the surrounding satisfactory voxels. Finally, spatial smoothing was applied within the CSF mask using the *3dBlurInMask* function from the AFNI software package(*44*), with a full width at half maximum value of 3 mm.

### Seed-based correlation analysis

Seed-based correlation analysis was performed separately in both OASIS cohorts. Regions of interest (ROIs) were defined in the template space by a neuroradiologist (A.N.) using ITK SNAP v3.8 (*45*). The selected ROIs included the premedullary cistern, cerebral aqueduct-4^th^ ventricle interface, foramina of Monro, left Sylvian fissure, and right Sylvian fissure. Mean MΨ values were extracted from these ROIs for each subject using the *fslmeants* function in FSL. Voxel-wise Pearson’s correlation maps were generated for each ROI using the *3dTcorr1D* function in AFNI. Voxelwise correlation values with false-discovery rate (FDR)-adjusted *p*-value < 0.01 were considered significant (corrected for five ROIs).

### Non-negative matrix factorization

We utilized NMF to identify the CSF flow patterns in which the CSF MΨ covaried consistently among participants. NMF is an unsupervised machine learning technique that decomposes a given non-negative matrix ***X*** into two non-negative matrices ***W*** (basis matrix) and ***H*** (coefficient matrix) such that ***X* ≈ *WH***. For the purpose of this study, the two-dimensional non-negative data matrix ***X*** was constructed by joining vectorized CSF MΨ maps in the template space (***X*** = 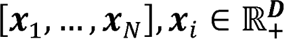, where ***D*** is the number of CSF voxels in the template space and ***N*** is the number of subjects). The resulting basis matrix 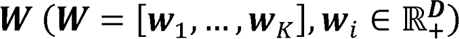 is composed of *K* columns, where *K* is the user-specified number of patterns. These columns represent the estimated CSF flow patterns. Each row in the coefficient matrix ***W*** represents a voxel, and the weights in the row indicate the relative contributions of that voxel to the CSF flow patterns. The matrix 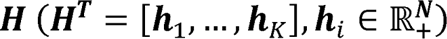 contains subject-specific coefficients for each CSF flow pattern, which indicate the contribution of each pattern in reconstructing the original CSF MΨ map. The implementation of NMF used in this study (https://github.com/asotiras/brainparts) enforces orthonormality constraints for the estimated covariation patterns (***W***^*T*^***W*** = ***I***, where ***I*** is the identity matrix) and projective constraints for their respective participant-specific coefficients (***H*** = ***W***^*T*^***X***) (*23*, *46*). The implementation of NMF has been discussed elsewhere in more details (*23*, *24*).

Consistent with prior studies using this technique (*22*–*24*), we ran multiple NMF solutions requesting 2–25 patterns to obtain a range of possible solutions for comparison. The optimal number of patterns (*K*) was selected based on the reconstruction error and stability of the solutions. The reconstruction error was calculated for each solution as the Frobenius norm between the data matrix and the NMF approximation. To determine the stability of our results (*22*–*24*), we performed a split-half reproducibility analysis with bootstrapping (with replacement) to examine the stability of NMF solutions across the range of possible resolutions (*47*–*49*). To ensure comparable age and sex distributions, the split-halves were generated using the *anticlust* package in R (*50*). We utilized two criteria to assess the stability of the NMF solutions: reproducibility, defined as the solution with the highest mean ARI across 20 split-half bootstraps, and reliability, defined as the solution with the lowest standard deviation of ARI across the bootstraps. ARI is a measure of set similarity that accounts for chance, with a value of 1 indicating a perfect match and a value close to 0 indicating a random partitioning (*28*). An ARI value greater than 0.75 is considered excellent (*51*).

### Replication analyses in the MICA and CDMD studies

*Low-b* dMRI data from the MICA dataset (b = 300 s/mm^2^) were corrected for Gibbs ring artifact, susceptibility-induced distortion using the FSL’s *topup* function and reverse phase-encoding b0 images. Data were additionally corrected for eddy-current-induced distortion and motion using FSL’s *eddy* function. Following skull-stripping with FSL’s Brain Extraction Tool (BET), the diffusion tensor model was applied to the preprocessed data to create MΨ maps. For the CDMD dataset (*36*), the available preprocessed dMRI data was used for further analysis. MΨ and FA maps were generated by fitting the diffusion tensor model to b = 50 s/mm^2^ dMRI data. Mean square displacement (MSD) maps were created by fitting the Laplacian-regularized mean apparent propagator (MAPL) MRI model (*52*) to the multi-shell dMRI data (b-values: 50, 350, 800 s/mm^2^) using *dipy* v1.5.0 (*53*). To assess the variability of MΨ, 100 sub-samples of b = 50 s/mm^2^ dMRI data were generated, each consisting of 12 volumes (with replacement). Coefficient of variance of MΨ values (standard deviation divided by mean) were calculated across the subsamples. In both MICA and CDMD datasets, the CWW masks from the OASIS-3 template space were nonlinearly registered to the individual MΨ images using the *antsRegistrationSyN.sh* script (*54*). Mean diffusion metric values from each CWW ROI were extracted in the native diffusion space.

### Volumetric analysis

T1-weighted images from the OASIS-3 dataset were segmented using FreeSurfer v5.3. (https://surfer.nmr.mgh.harvard.edu/) (*55*). The image segmentations were reviewed by a trained lab member of the OASIS project (*34*). TkMedit (part of FreeSurfer; http://freesurfer.net/fswiki/TkMedit) was used to revise images that failed the quality control. The revised images were rerun through the FreeSurfer pipeline. Total intracranial volume, brain parenchymal fraction (brain parenchymal volume divided by total intracranial volume), and ventricular volumes (normalized by total intracranial volume) were extracted for further analyses.

### Cerebral blood flow quantification

Cerebral blood flow (CBF) quantification was performed using the *oxford_asl* package v4.0.27, part of the BASIL toolbox within FSL (*39*, *56*). This procedure involved motion correction, automated spatial regularization, label-control subtraction, relative CBF quantification, and conversion of relative CBF to absolute physiological units (ml/100 g/min) using the M0 image (*56*). Model parameters were selected based on the ASL Consensus Paper for a magnetic field strength of 3 T(*57*): arterial blood longitudinal relaxation time (T_1b_), 1650 ms; brain tissue longitudinal relaxation time (T_1t_), 1300 ms; inversion efficiency (α), 0.98. The TI_2_ value was adjusted slice by slice with an inter-slice acquisition time difference of 46 ms. A gray matter mask (gray matter partial volume effect > 0.5; derived from the *fsl_anat* script) was created in the ASL native space and used to extract mean gray matter CBF values.

### Neuroradiological evaluation

In the OASIS study, aberrant CSF flow patterns were defined as either (i) one or more CWW with an MΨ deviating more than 3-standard deviations from the mean, or (ii) two or more CWWs with MΨ values deviating more than 2-standard deviations from the mean. In order to evaluate any related incidental findings and/or potential underlying etiologies of these aberrant CSF flow patterns, available MRI images of OASIS study participants displaying such patterns were evaluated by a neuroradiologist (A.N.).

### Statistical analyses

All region-of-interest statistical analyses were performed with R v4.1.2. (http://www.r-project.org/). To determine the study-specific effect sizes (standardized β) of age, sex, and brain parenchymal fractions (BPF) on CWW MΨ values, general linear models were fitted separately for each of the two OASIS cohorts (Model 1: CWW_MΨ ∼ age + sex + BPF). Effects of ventricular volumes and CBF were estimated while accounting for the effects of age, sex, and BPF (Model 2: CWW_MΨ ∼ age + sex + BPF + [ventricular volume/CBF]). Results of the two OASIS cohorts were aggregated using linear mixed-effects models (*lmer* function, part of the *lme4* package) with a random intercept for the cohort (Mixed Model: CWW_MΨ ∼ age + sex + BPF + (1|Cohort)). The p-values of the linear mixed models and CBF general linear models were adjusted for multiple testing using the false discovery rate (FDR). FDR-corrected p-values lower than 0.05 were considered significant.

## Acknowledgments

A.S. was partially supported by the National Institutes of Health (NIH) award R01AG067103. The facilities of the Washington University Center for High Performance Computing (CHPC) partially funded by NIH grants 1S10RR022984-01A1 and 1S10OD018091-01, were used for computational analyses.

## Notes

### Competing Interest Statement

The authors have declared no competing interest.

