## Supplementary figures and images for "The Spatial Patterns and Determinants of Cerebrospinal Fluid Circulation in the Human Brain"

### Figure S1

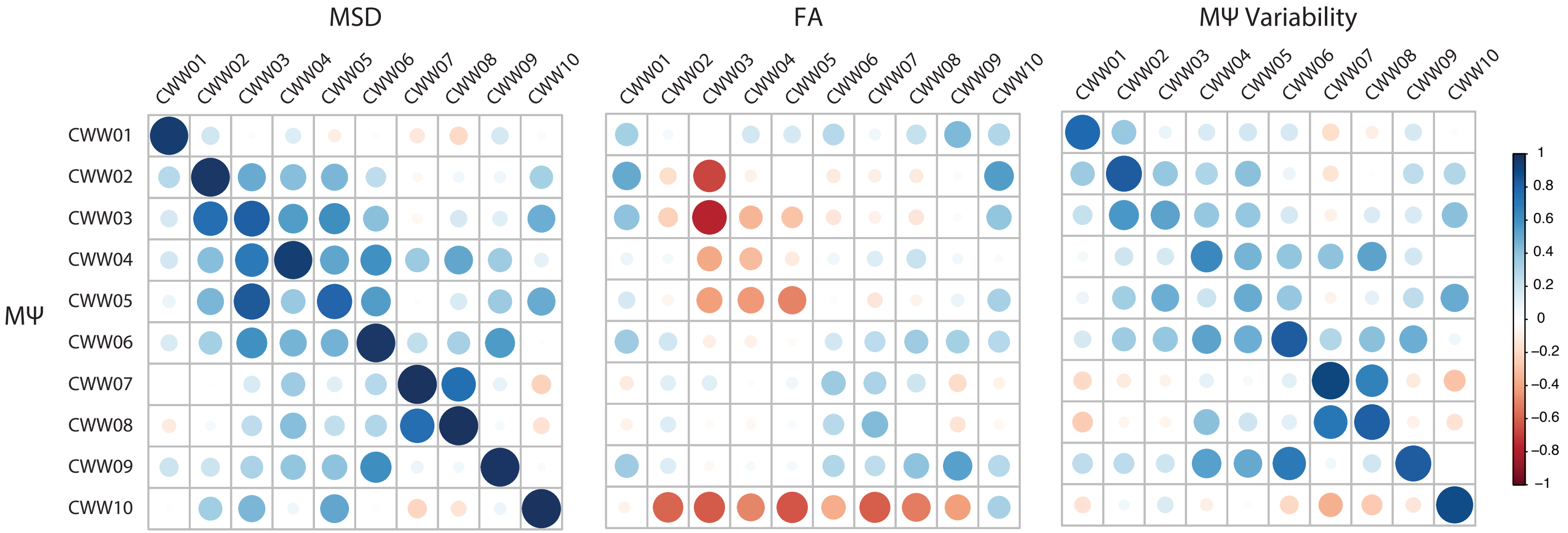
